# Multiplexed embryo profiling links cellular state to zygotic genome activation in single cells

**DOI:** 10.1101/2025.10.27.684723

**Authors:** Max Hess, Marvin F. Wyss, Edlyn Wu, Joel Lüthi, Chiara Rebagliati, Nadine L. Vastenhouw, Darren Gilmour, Shayan Shamipour, Lucas Pelkmans

## Abstract

Multicellular self-organization relies on complex interactions across multiple length scales, shaping the transcriptional activity and molecular phenotype of individual cells. Despite advances in spatial omics technologies, mapping protein state composition at high spatial resolution to capture the molecular phenotype across whole-mount structures has remained challenging. Here, we introduce high-throughput 3D iterative indirect immunofluorescence imaging (**3D-4i**) for the simultaneous detection of multiple proteins and protein states across the entire embryo, and a 3D-dedicated computer vision pipeline for quantifying morphological and molecular properties across subcellular, single-cell, and whole-embryo scales in hundreds of samples. Applying this pipeline to early zebrafish embryos undergoing mid-blastula transition, we determine the cell cycle phase for each cell across the embryo, allowing us to uncover the spatiotemporal dynamics by which global meta-synchronous mitotic waves transition to cell cycle desynchronization. Using statistical analysis, we find that cell cycle phase is the major source of variability in transcription within a division cycle, and combining this with analysis of key transcription factors and chromatin modifier state enables accurate prediction of transcriptional output during zygotic genome activation in individual cells. Collectively, our results demonstrate the strength of 3D-4i in quantifying multimodal effects across spatiotemporal scales and show how it can be utilized to unravel the complex contribution of causal factors that collectively drive multicellular self-organization.

## Introduction

A crucial stage in early embryonic development is marked by the transition from maternally provided transcripts to zygotic genome activation (ZGA). In many species, the major wave of ZGA coincides with the lengthening of cell cycles mediated by the introduction of gap phases into the early rapid divisions^1^ (known as mid-blastula transition, MBT). While multiple factors have been causally related with ZGA, including slowing down of the cell cycle during MBT^2,3^, titration of replication machinery^4,5^, dilution and post-translational modification of maternally supplied histones^6,7^, and the action of pluripotency factors such as Pou5f3 and Nanog^6,8–10^, ZGA is thought to be imprecisely controlled in single cells due to its highly heterogeneous onset^11^. We here explore the possibility that the heterogeneous nature of ZGA reflects the interplay of multiple causal effects emerging at different spatiotemporal scales, which collectively ensure a predictable activation of the zygotic genome at different timepoints in individual cells to generate reproducible patterns of transcriptional activation in early embryos. Identifying sources of cell-to-cell variability requires the ability to quantify multiple contributing effects simultaneously in a large number of individual cells within their native spatial context^12–14^.

To achieve this in a developmental system, we leveraged the accessibility, abundance, and optical transparency of zebrafish embryos and established a high-throughput approach that combines 3D *in toto* imaging of whole-mount structures with iterative indirect immunofluorescence imaging^15^ (3D-4i). Using this technology, we imaged multiple protein markers across all cells in the entire zebrafish embryo at subcellular resolution. Additionally, we implemented a comprehensive 3D image analysis pipeline to extract features across whole-embryo, single-cell, and subcellular scales for downstream analysis. By applying 3D-4i to a large set of zebrafish embryos undergoing MBT, we capture the multivariate and multiscale nature of processes involved in ZGA at single-cell resolution, and show that these measurements allow the training of models that can accurately predict the transcriptional activity of individual cells during ZGA, identifying a major role for position along the cell cycle, in combination with singlet-cell levels of ZGA-associated factors, particularly Pou5f3, PCNA and H3K27ac. This underscores the strength of embryo multiplexing in disentangling the contributions of causal factors to complex processes in development.

## Results

### Establishing 3D iterative indirect immunofluorescence imaging in zebrafish embryos

To achieve multiplexed molecular profiling in whole-mount embryos, we adapted the 4i technology, previously applied to mammalian cells grown in culture and 2D tissue sections^13,15,16^, enabling homogeneous immunofluorescence staining, 3D volumetric imaging and efficient antibody elution, thereby facilitating the sequential addition of new antibody sets (3D-4i, Fig. 1a, Materials and Methods). A critical step in establishing this workflow was robust sample immobilization, which enabled precise computational alignment of embryos across imaging cycles (Fig. S1). To achieve this, we mounted deyolked zebrafish embryos on Fibronectin/Poly-D-Lysine-coated 96-well plates using UV photocrosslinking. This method, combined with a gentle liquid handling system for dispensing and aspirating solutions, minimized embryo displacement and loss, while maintaining embryo morphology during multiplexing cycles. In addition, homogeneous antibody staining of high-abundance targets across the 3D embryo volume was achieved by directly labeling primary antibodies with fluorescently tagged Fab fragments. Finally, embryos were optically cleared using refractive index-matching solution, allowing volumetric imaging through tissue depths exceeding 250 µm (Materials and Methods). To obtain a dataset for measuring transcriptional activity and its associated factors in individual cells, we performed 3D-4i on 212 early zebrafish embryos, encompassing a total of 458,826 individual cells, collected between 2 and 4 hours post-fertilization (hpf, corresponding to the 7^th^ to 12^th^ division cycles) at 15-minute intervals, the developmental time window during which the major wave of ZGA occurs (Fig. 1b). Assuming uniform sampling of embryos, this dataset corresponds to a subminute temporal resolution during this time (120 min / 212 embryos ∼ 34 sec). Each embryo was imaged at 20x magnification using automated spinning-disk confocal microscopy across four imaging rounds, generating a volumetric dataset that includes DAPI to quantify DNA content (also used for image registration and nuclear segmentation) and ten-plex protein stainings capturing key molecular markers of the ZGA process. These comprised indicators of cell cycle progression (PCNA, an S phase marker^17^, and phosphorylated H3, pH3, as a marker for mitosis^18^), histone abundance and modifications (H2B and K27-acetylated H3, H3K27ac) reflecting chromatin state, pluripotency transcription factors initiating the zygotic developmental program (Pou5f3 and Nanog), an mRNA nuclear export factor (Alyref), the initiating and elongating forms of RNA Polymerase II (Pol-II-S5p and Pol-II-S2p, respectively) as reporters of transcriptional activity^19^, and finally β-catenin for cell and embryo segmentation (Fig. 1c-d). Importantly, embryos of different timepoints were mixed and distributed across 30 wells (∼7-9 embryos per well) to reduce technical error due to well-to-well variations.

**Fig. 1:**
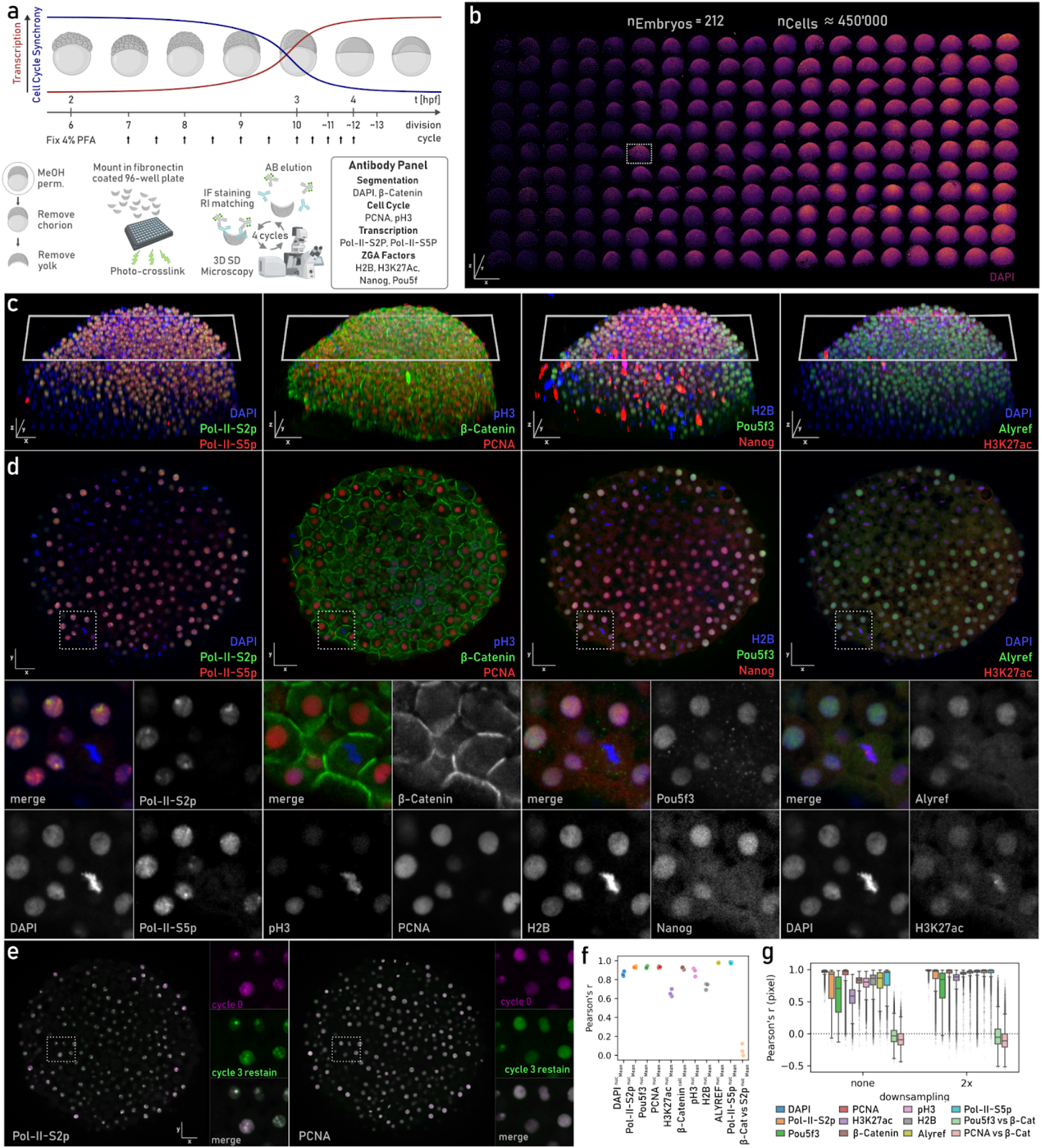
Establishing 3D iterative indirect immunofluorescence imaging (3D-4i) in zebrafish embryos. a. Schematic of zebrafish developmental timepoints used in the multiplexing experiment (top), 3D-4i workflow (bottom left), and antibody panel used in the experiment (bottom right). MeOH perm: Methanol permeabilisation. SD: Spinning Disc. RI: Refractive Index. IF: Immunofluorescence. AB: Antibody. For further details, see Materials and Methods section. b. 3D overview of embryos imaged in the multiplexing experiment, ordered from left to right by total nuclei count. Images show the DAPI stain labeling DNA. The ROI demarcates a representative embryo shown in panels c and d. x, y, z scale bars 650 µm. c. 3D view of a representative embryo (demarcated in panel b), stained with Pol-II-S2p, Pol-II-S5p, β-Catenin, PCNA, pH3, Pou5f3, H2B, Nanog, Alyref, H3K27ac, and DAPI across four sequential immunofluorescence imaging cycles. The white rectangles demarcate the 2D slice used for visualization in panel d. x, y, z scale bars 50 µm. d. 2D cross-section of the embryo shown in panel c along the indicated plane, with zoom-in views shown below. The dashed rectangles demarcate the ROI used for zoom-in. x, y scale bars 50 µm. e. Stainings of Pol-II-S2p and PCNA in acquisition cycles 0 and 3 overlaid on each other, with zoom-in views shown on the right. The dashed rectangles demarcate the ROI used for zoom-in. x, y scale bars 50 µm. f. Pearson’s r for mean nuclear intensities (cellular for β-Catenin) between same stains repeated across cycles. β-Catenin (β-Cat) vs Pol-II-S2p (S2p) serves as negative control. g. Single-pixel correlations between same stains repeated across cycles, at full image resolution (1 pixel: 0.325 µm, left: none) or after 2× down-sampling (1 pixel: 0.650 µm, right, 2x). β-Cat vs PCNA or Pou5f3 serve as negative controls.

To assess the reproducibility of staining intensities across imaging cycles, we re-stained markers from the first three cycles in the final cycle, using three replicate wells each, and conducted a correlation analysis between the original and re-stained images (Fig. 1e). Mean nuclear (or cellular for β-catenin) intensities showed high single-cell correlations between the same stains across different imaging cycles and low correlations between unrelated stains (β-catenin and Pol-II-S2p) (Fig. 1f). High single-pixel correlations between the same stains, either in the full resolution or in two times down-sampled images further validated the reproducibility of the stains in subcellular scale (Fig. 1g). We noticed a slight accumulation of background signal over cycles, resulting in reduced single-pixel correlations of targets with low signal-to-noise ratio (H3K27ac and Pou5f3). Collectively, these results demonstrate that iterative staining in embryos is highly reproducible and preserves sample integrity also at small length scales, establishing 3D-4i as a high-throughput, whole-mount multiplexed immunofluorescence approach that enables simultaneous investigation of spatiotemporal patterns of multiple protein states in hundreds of zebrafish embryos.

### Developing an integrated image analysis pipeline for multiplexed volumetric datasets

To extract information from these multiplexed volumetric datasets in a systematic and reproducible manner, we next generated a comprehensive image analysis pipeline, encompassing image file compression, multiscale image pyramid generation, intensity correction accounting for various technical aberrations (as detailed below), image registration across staining cycles, and nucleus, cytoplasm, cell and embryo segmentation through the integration of multiple established toolboxes^20–28^ (Fig. 2a-g; S1-S2 see Materials and Methods).

**Fig. 2:**
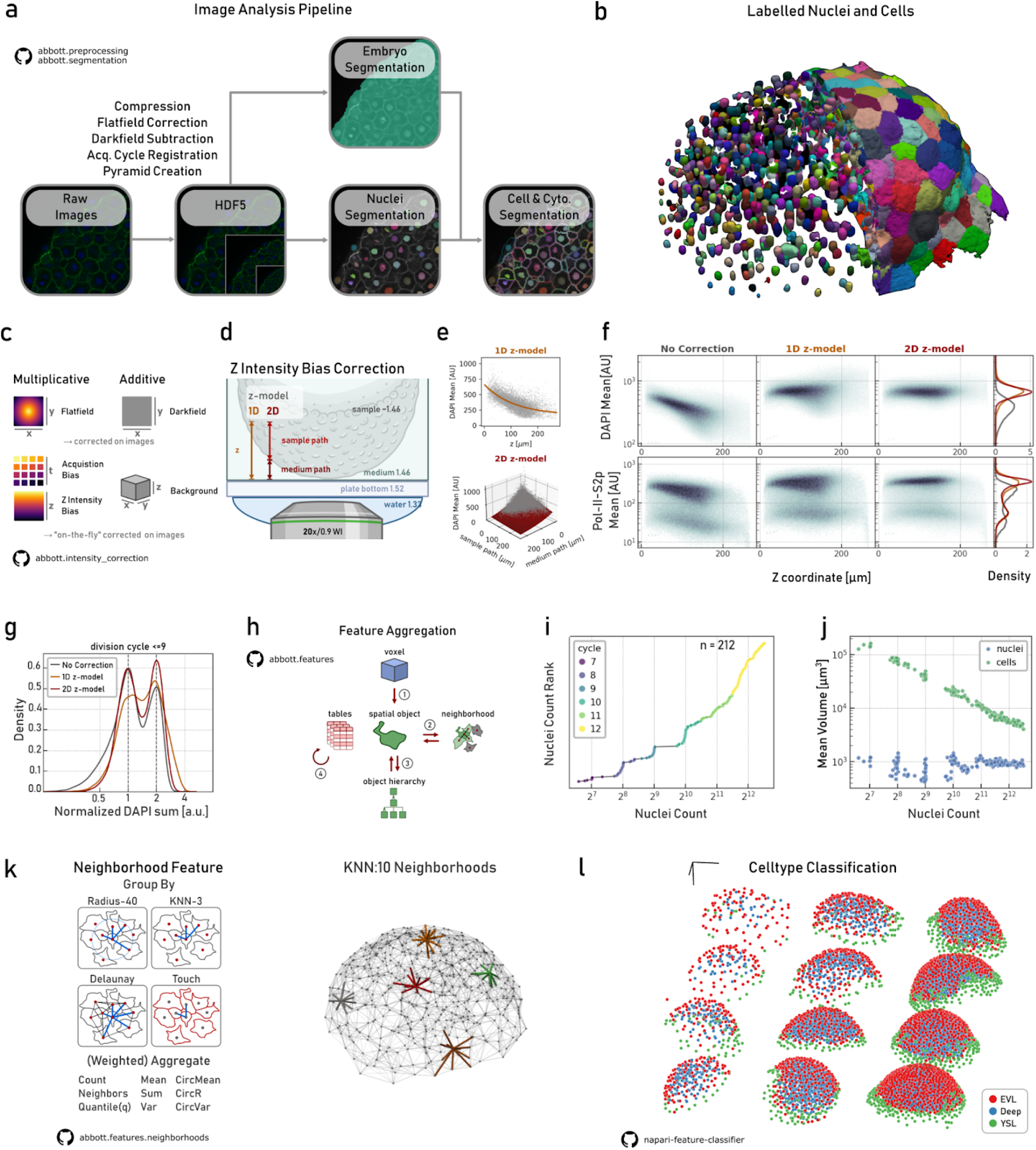
Developing an integrated image analysis pipeline for multiplexed volumetric datasets. a. Image-analysis pipeline including conversion into multiscale image pyramids, intensity correction, image registration between acquisition (Acq.) cycles and object (embryo, nuclei, cell and cytoplasm) segmentation. b. 3D view of a representative embryo with nuclear segmentation shown on the left half and cell segmentation on the right half. c. Series of intensity corrections applied directly to images or corrected on the fly (See Materials and Methods). d. Schematic of exemplary cell position relative to the objective, showing the light path through the plate bottom, medium, and sample, with their approximate refractive indices. The single-exponential 1D z-decay model only considers the total light path (medium and sample paths sum) and the double-exponential 2D z-decay model considers separate decays for each path. e. Estimation of mean DAPI signal decay using 1D and 2D z-models. f. Comparison of 1D and 2D z-decay models for mean DAPI and Pol-II-S2p intensity distributions. g. The density plot on comparison of 1D and 2D z-decay models with uncorrected data for summed DAPI intensity of embryos in cycles 7-9. Data normalized to the first peak. h. Logical data model for feature extraction across scales/orders, dividing the process into two steps of label aggregation (across voxels (1), neighborhoods (2), object hierarchy (3) or tables (4)) and computation of summary statistics on aggregated labels. i. Embryo staging based on total nuclear counts. j. Mean nuclear and cellular volumes plotted against total nuclear counts per embryo. k. Left: various statistical features (such as mean, sum, etc) can be aggregated across different neighborhoods (fixed radius, touch, k-nearest neighbors (KNN) and Delaunay, Materials and Methods) to achieve higher scale measurements. Right: Example for KNN neighborhoods with k=10 in an embryo. Neighbors of exemplary cells shown in colors and thicker lines. l. Results of a random forest classifier trained with the napari feature-classifier plugin to identify three cell types present during early zebrafish development: EVL, YSL and deep cells (Deep).

Central to image analysis is to ensure a linear relationship between measured signal intensities and actual fluorophore concentrations. To achieve this, we applied a series of signal corrections either directly to the images or during downstream analysis. First, we applied darkfield and flatfield corrections to the images, which account for thermal noise, camera offset and uneven illumination^28^. Second, we measured background signals by the increase in cytoplasmic signal for each stain, which particularly accumulated in later cycles due to incomplete antibody elution, and corrected this on the extracted intensity features. Third, we estimated for each stain exponential intensity decays caused by fluorophore quenching in the refractive index-matching solution over time (acquisition bias) and along the imaging axis (z-decay bias, Fig. S2) due to refractive index mismatches and inhomogeneous scattering in the tissue, and corrected these on the image volumes during post-processing (Fig. 2c; Materials and Methods). We used a single exponential model, applied to the mean intensity decay of stains over acquisition time, to account for acquisition bias. For addressing z-decay bias, we tested two models: (a) a single exponential model, which accounts for intensity decay based on the total light path from the objective to the focal plane (1D z-model), and (b) a double exponential model, that separates the light path into the medium and sample distances, accounting for their distinct scattering properties and intensity decay rates (2D z-model) (Fig. 2d).

To evaluate the performance of these correction models, we used the mean and sum nuclear DAPI intensity distributions as ground truth. For mean nuclear DAPI intensity, which is expected to be uniform throughout the early embryo, we noted that both z-decay correction models flattened the observed mean intensity decay over the tissue depth, with the 2D z-model further improving the homogeneity of intensity variance along the z-axis. The 2D z-decay model also accounted for intensity decay in other stains, such as Pol-II-S2p, used as a marker for transcriptional output (Fig. 2e-f). For sum nuclear DAPI intensity, we expect a bimodal distribution, with peaks separated by a factor of two, corresponding to nuclei with two or four DNA copies, based on the cell cycle stage. The 2D model more clearly resolved these peaks and reduced variability around the expected values compared to both the 1D model and uncorrected data, highlighting the necessity and strength of this approach in correcting intensity decays (Fig. 2g and S2).

After intensity corrections and segmentation of spatial objects (e.g., nuclei and cells, see Materials and Methods), we focused on extracting biologically meaningful features from these high-dimensional data comprising approximately 10^9^ volumetric pixels (voxels) per embryo, across 212 embryos, each with ten-plex protein stainings. To streamline this process, we developed a logical data model that separates feature extraction into two steps of label aggregation and the computation of summary statistical functions on the aggregated labels. In this framework, object measurements such as nuclear volume or mean intensity are acquired by applying statistical functions to aggregated voxels within the same label objects (e.g., nuclei labels). Similarly, we can aggregate features for a given object type, over a series of spatial neighborhoods or across different levels of object hierarchy to derive features at higher length scales/orders. Finally, feature embeddings or cell type classifications can be viewed as aggregating existing features to generate higher-level properties (Fig. 2h).

For instance, to identify the developmental stage of embryos, we can aggregate nuclei objects at the embryo hierarchy level and sum their total counts. As expected, we find that, up to the 10^th^ cell cycle, the nuclei count clustered around powers of two, reflecting the rapid and meta-synchronous nature of early embryonic division cycles. However, as synchronicity is lost in later cycles^17,29^, a more gradual increase in nuclei count between embryos is observed (Fig. 2i). Furthermore, plotting mean nuclear and cellular volumes against the number of nuclei per embryo revealed a halving of cell volume with each division cycle, while nuclei volume remained relatively constant (Fig. 2j). Similarly, local cell density in the embryos can be measured, for instance by aggregating cell count within a 50 *μ*m radius neighborhood, revealing that density varies locally, and is typically higher in the center than in the periphery of the embryo (Fig. 2k and S3a). Leveraging this logical data model, features such as position, nuclear morphology and mean intensity of stains were aggregated to train a random forest classifier that distinguishes fluorescent debris from actual nuclei and enables unbiased identification of the three cell types observed during early zebrafish development: the enveloping layer cells (EVL), yolk syncytial layer (YSL), and deep cells giving rise to embryo proper (Fig. 2l and S3b, Materials and Methods). The classifier’s predictions were then used to remove debris and exclude YSL cells from downstream analysis, as they often exhibit distinct nuclear morphologies in comparison to deep and EVL cells^30^. Collectively, this logical data model serves as a unifying framework for feature aggregation, enabling the integration of new features and aggregation modes. Together with the developed image analysis pipeline, this supports robust measurements across multiple spatial and hierarchical scales.

### Inferring cell cycle phase from multiplexed imaging of fixed embryos reveals spatiotemporal dynamics of cell cycle desynchronization during mid-blastula transition

CCP is a known source of variability in transcriptional activity during ZGA^1–3,10,11,31^, thus we developed methods to reliably infer CCP in fixed samples, enabling analysis of the impact of early cell cycles on transcriptional output. Since many cellular and nuclear features change throughout CCP, we hypothesized that a randomly selected population of cells should organize along a circular manifold in a high-dimensional feature space defined by cell cycle-related attributes such as nuclei morphology, DAPI and PCNA intensities^32^. This manifold should allow the estimation of each cell’s position along the cell cycle trajectory, providing a systematic approach for the analysis of cell cycle dynamics from fixed images. To this end, cells were embedded in lower-dimensional spaces using Uniform Manifold Approximation and Projection (UMAP^33^), following pre-processing steps such as data subsampling to address class imbalances across division cycles and feature selection/normalizations (Fig.3a, Materials and Methods). As expected, cells pooled from all timepoints organize along a circular manifold in a 3-dimensional UMAP embedding of cell cycle features. While the first two UMAP components capture cell cycle progression, the third axis largely corresponds with changes related to consecutive division cycles (Fig. 3b). A principal circle was then fitted to the embedded nuclei of each division cycle to model the continuous and circular nature of cell cycle progression and correctly map 634 manually annotated nuclei to their corresponding cell cycle phase (Fig. 3b-b’, Materials and Methods). The CCP value for each cell, which we bounded between 0 and 2π, was then defined from the closest point on the principal circle of its respective division cycle (Materials and Methods). Finally, because distinct cell cycle phases have different durations, resulting in a non-uniform distribution of cells along the cell cycle (Fig. 3b), we normalized the CCP density (hereafter termed CCP), resulting in a uniform distribution that reflects the progression of cells through the cell cycle (Fig. 3c).

**Fig. 3:**
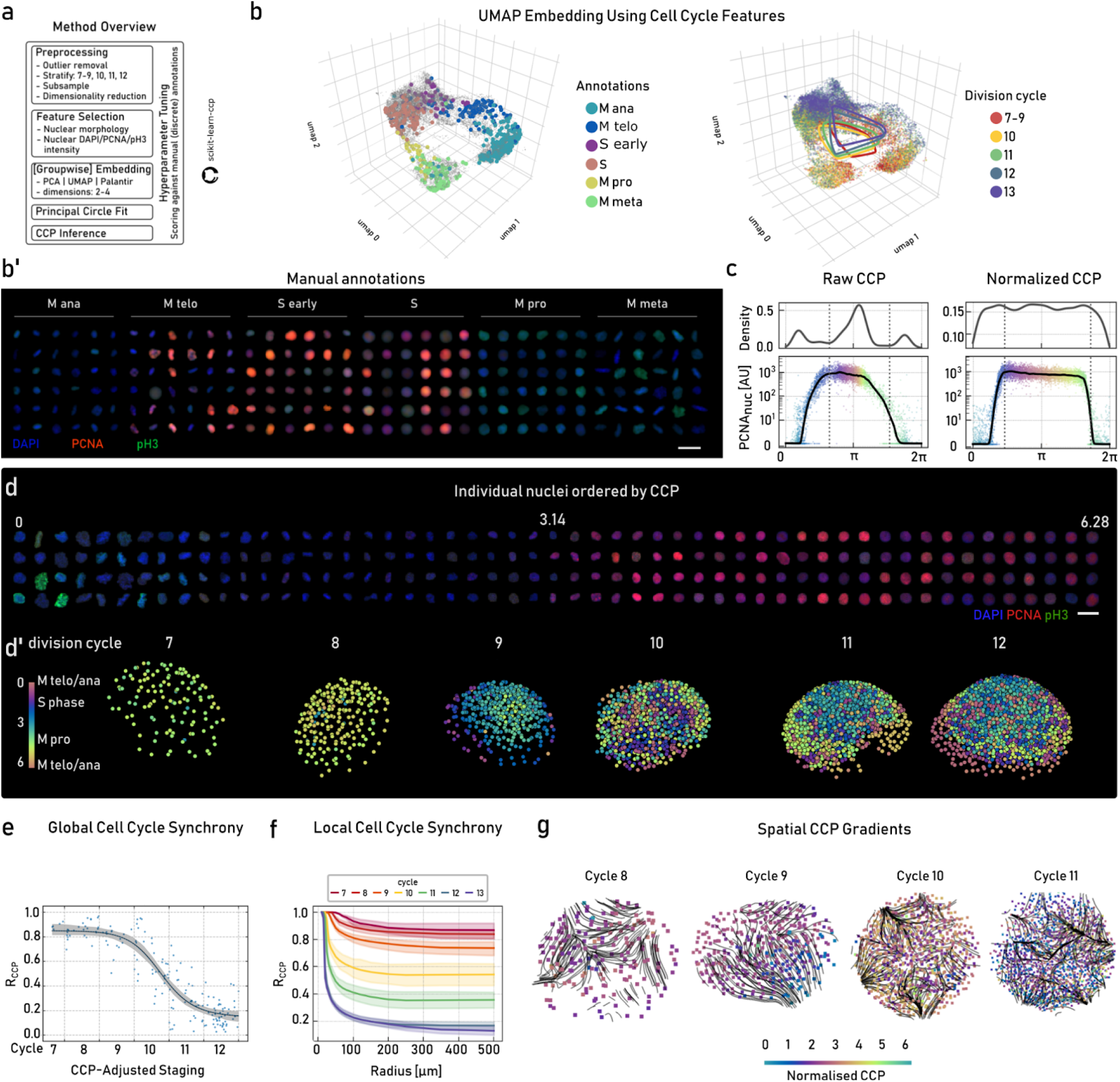
Inferring cell cycle phase from multiplexed imaging of fixed embryos reveals spatiotemporal dynamics of cell cycle desynchronization during mid-blastula transition. a. Method overview for pre-processing, single-cell embedding into a low-dimensional space using cell cycle-related features, principal circle fitting in feature space and CCP inference (Materials and Methods). b. Left: UMAP embedding of cell cycle features overlaid with manual cell cycle annotations. Right: Principal circles fitted to feature space of cells from division cycles 7–9, 10, 11, 12, and 13 separately. b’. Representative nuclei annotations for six cell cycle phases visualized using DAPI, PCNA, and pH3 markers. Scale bar 20 μm. c. Density plot and mean nuclear PCNA intensity of 10,000 cells randomly sampled across division cycle 11, plotted against raw CCP (left) or normalized CCP (right, hereafter CCP) after inference and quantile-based normalization. Points are colored by normalized CCP, illustrating the monotonic mapping introduced by normalization. Black line: LOWESS fit highlighting observed dynamics. Dotted vertical lines correspond to M anaphase/telophase, S, and M pro/meta phases. d. Exemplary nuclei ordered by inferred CCP and visualized with DAPI, PCNA, and pH3 markers. Scale bar 20 μm. d’. Representative embryos from division cycles 7 to 12 with nuclei shown as point clouds and colored by their inferred CCP. e. Global cell-cycle synchrony (R) plotted against CCP-adjusted staging, fitted with a Hill function (Materials and Methods). The solid line is the bootstrap-mean fit and the shaded region shows the pointwise 95% bootstrapped confidence interval. f. CCP synchrony measured over increasing radii across division cycles 7-13. Solid lines indicate group medians for the corresponding division cycle and the shaded regions mark 95% bootstrapped confidence intervals. g. Spatial CCP gradients for 4 representative embryos from division cycles 8 to 11. Streamlines correspond to the integrated gradient field starting from 500 randomly sampled sources within the convex hull of nuclei positions. Nuclei colors denote the inferred CCP.

Having inferred the CCP values for all cells within each embryo (excluding YSL nuclei), we refined embryo staging based on the circular mean CCP in combination with nuclear count (CCP-adjusted staging, Fig. S4, Materials and Methods). When visualizing the CCP of all cells in their spatial contexts, we observed that in early embryos (division cycles 7 to 9), cells exhibit similar CCP values (meta-synchronous), whereas with increasing developmental time, the cell cycle becomes asynchronous across the embryo (division cycles 10-12, Fig. 3d-d’). To quantify the dynamics of the transition from meta-synchrony to asynchrony, we measured the circular synchrony of the CCP (R) for all cells within each embryo (Materials and Methods), where R = 1 indicates perfect synchrony, and R = 0 corresponds to complete asynchrony. This analysis revealed that up to cycle 10, average CCP synchrony remains high (above 0.7), but drops sharply at cycle 10 (fitted Hill coefficient of ≈ 22, Materials and Methods) and approaches near-complete asynchrony (∼0.1) at cycles 11 and 12 (Fig. 3e). This abrupt increase in CCP heterogeneity at division cycle 10 coincided with a switch-like rise in transcriptional activity, as indicated by measurements of Pol-II-S2p foci volume (Fig. S5), which carry the majority of nascent RNAs^3^, pointing at the potential causal link between CCP dynamics and zygotic genome activation (see Fig. 4). In addition, we analyzed whether there is a spatial component to cell cycle desynchronization by quantifying CCP synchrony between cells positioned at increasing distance from each other (from 0 to 500 μm reaching the embryo boundaries). This analysis demonstrated a half maximum synchrony at ∼100 μm in cycle 7 embryos that steadily decreased to ∼25 μm by cycles 12 and 13, corresponding to approximately one cell diameter (Fig. 3f), suggesting that a gradual decrease in spatial synchrony might underlie the observed rapid temporal transition in global CCP synchrony. In line with this, calculating spatial gradients of mitotic progression using the inferred CCP values (Materials and Methods), revealed coherent pseudo-waves in cycles 8 and 9, and more chaotic local patterns in cycles 10-11 (Fig. 3g), as previously reported based on live-imaging experiments^17,29^. Altogether, by integrating machine learning algorithms and statistical approaches, we inferred CCP at the single-cell resolution, enabling quantification of the spatiotemporal dynamics of cell cycle transitions from meta-synchrony to asynchrony and establishing the foundation for assessing the role of CCP in regulating transcriptional activity.

**Fig. 4:**
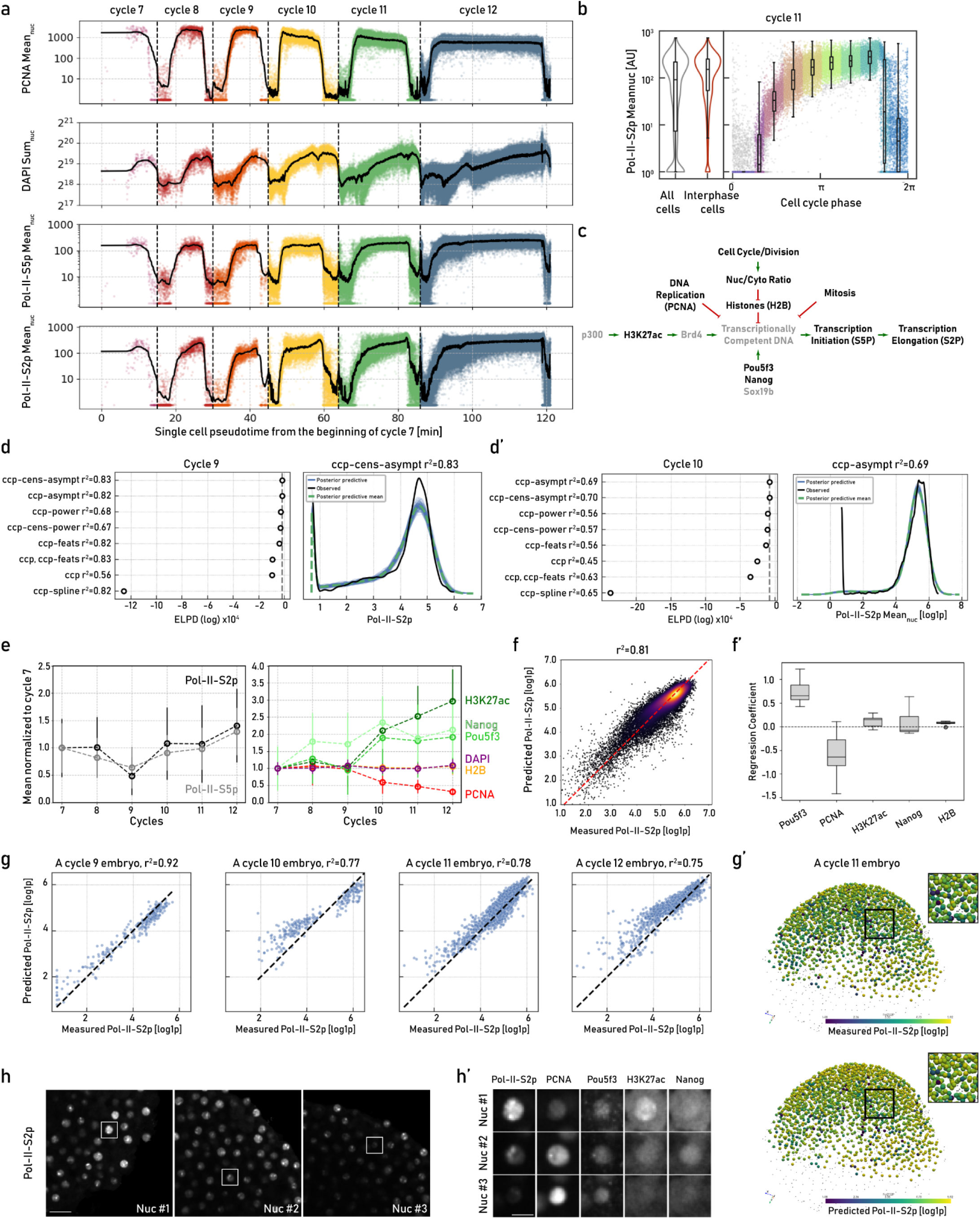
Multiplexed embryo profiling links cellular state to zygotic genome activation in single cells. a. Mean PCNA, Pol-II-S5p and Pol-II-S2p and sum DAPI nuclear intensities across division cycles 7 to 12. Sold lines demarcate Gaussian smoothing of points and dashed lines and color changes indicate cycle transitions. b. Left: Mean nuclear intensities of Pol-II-S2p distributions for all (gray) and interphase (red) cells during division cycle 11. Right: Same data plotted against their inferred CCP. Data points were divided into nine bins across CCP (shown with distinct colors, six equally sized interphase bins and three mitotic bins). The distribution within each bin is shown with its overlaying boxplot. c. Causal interactions known to contribute to transcriptional output^10,56^. Green/red arrows indicate activating/repressive interactions. Gray components represent known regulators not measured in this dataset. d-d’. Left: Model comparisons using CCP or features used for inferring CCP (CCP features, see Materials and Methods) as linear or non-linear predictors of transcription output measured by natural logarithm of mean nuclear Pol-II-S2p for cycles 9 and 10. Models are ranked based on their performance on held-out data using expected log pointwise predictive density ELPD (improving from left to right). R^2^ demarcates explained variance for each model. Asymptotic (asympt), power and spline non-linear regression models of CCP were tested. Linear models include CCP or features used for CCP inference (ccp-feats). For the asymptotic and power regressions, additional models were tested where data was clipped at 100 gray values, censoring data points with low signal-to-noise ratios (ccp-cens-asympt, ccp-cens-power, Materials and Methods). Right: Posterior predictive distributions (blue) and their mean (green) of the best-performing model compared to the observed data (black). Panels d and d’ correspond to data and models from cycles 9 and 10, respectively. e. Mean interphase changes of transcriptional reporters Pol-II-S5p and Pol-II-S2p (left) and ZGA-associated factors H3K27ac, Pou5f3, Nanog, H2B, and PCNA together with DAPI (right) across division cycles 7 to 12. f. Measured versus predicted Pol-II-S2p levels (natural log-transformed) for all interphase cells with CCP held constant. S phase was divided into six equally sized strata and a multilinear regression was applied to each bin (Materials and Methods). Color code demarcates point density and red dashed line is the line of equality. f’. Regression coefficients for each measured ZGA-associated factor across six interphase bins. g. Measured versus predicted Pol-II-S2p levels (natural log-transformed) for three single held-out embryos using the best-performing model for the corresponding cycles 9 to 12. Black lines demarcate the lines of equality. g’. Measured (top) and predicted (bottom) mean nuclear Pol-II-S2p (natural log-transformed) in a cycle 11 embryo. Gray dots denote YSL nuclei, mitotic cells, segmentation artifacts, or intensity outliers in any staining that were excluded from the analysis. Squared boxes demarcate the zoom-in views shown in top-right panels. h. 2D sections of a representative embryo from cycle 11 (same embryo as shown in g’) stained with Pol-II-S2p antibody. White boxes indicate the exemplary nuclei (nuc) shown in h’. Scale bar 25 µm. h’. Exemplary nuclei, indicated in panel h, stained with Pol-II-S2p, PCNA, Pou5f3, H3K27ac and Nanog antibodies. Scale bar 10 µm.

### Multiplexed embryo profiling links cellular state to zygotic genome activation in single cells

We next investigated whether spatially resolved measurements that capture multiple features across biological scales provide contextual information that explains the heterogeneity observed in transcriptional activity of individual cells during zygotic genome activation. Because CCP is a major source of variability in cellular states of mammalian cells grown in culture^15^ and becomes abruptly asynchronous in division cycle 10 embryos coinciding with ZGA onset (Fig. 3e and S5), we visualized variation of the measured features in our multiplexing experiment across CCP. To this end, we generated a pseudotemporal axis of cell cycle progression from division cycle 7 to 12 by mapping individual cells onto developmental time, using their embryonic stage and inferred CCP scaled by division cycle durations obtained from live-imaging experiments^29^ (Fig. 4a and S6a). Examining DAPI sum intensity demonstrated that DNA content dropped immediately after chromosome segregation and doubled during S phase, while mean PCNA intensity showed rapid transitions between high values in S phase and low values in M phase, as expected^17^ (Fig. 4a).

Plotting mean nuclear intensity of Pol-II-S5p and Pol-II-S2p along this axis as a proxy for average transcriptional activity revealed two types of dynamics: A slow, gradual increase across successive division cycles, as well as variations within each cell cycle, characterized by low activity of RNA Pol-II during mitosis followed by a sharp increase in activity during S phase, eventually reaching a plateau (Fig. 4a). Consistent with this cell cycle-dependent dynamics, variation in transcriptional activity between individual cells of a given division cycle was markedly reduced when considering interphase cells only or cells belonging to similar CCP (Fig. 4b). To investigate the dependency between CCP and transcriptional output, we implemented a Bayesian regression framework to predict zygotic transcriptional activity in single cells, as measured by mean nuclear Pol-II-S2p during interphase, using the inferred CCP of each cell with either linear or non-linear functions (Materials and Methods). Model performance was assessed by explained variance (R^2^) and prediction accuracy was evaluated on held-out data, revealing that the relationship was best described by a non-linear asymptotic regression model (ccp-asympt, Fig. 4d-d’ and S6b). Using our modeling framework, we found that CCP alone is a strong predictor of transcriptional activity, explaining between 65-85% of the variance across individual cells in division cycles 8-12 (Fig. 4d-d’ and S6b)^11^.

To study the variance in transcriptional activity not explained by CCP, we next quantified the abundance of the proteins associated with ZGA measured in our multiplexed dataset (Fig. 4c). At the embryo-averaged level, we observed a pronounced decrease in PCNA as division cycles progressed, concomitant with a rise in transcription initiation and elongation, consistent with previous model proposing that titration of replication machinery is required for transcriptional activity^4,5,34^ (Fig. 4e). In contrast, the pluripotency transcription factors Nanog and Pou5f3, together with the active chromatin mark H3K27ac increased markedly from division cycle 10 onwards (Fig. 4e), indicating that the combined chromatin and transcription factor landscape progressively acquires transcriptional competence during MBT^6–8,35^. Importantly, at the single-cell level, these markers displayed substantial heterogeneity, which paralleled the increasing heterogeneity in transcriptional output observed during later cycles (Fig. 4e and S5a’). To disentangle CCP-dependent from CCP-independent contributions to transcription, we performed multilinear regression modeling using ZGA factors (Pou5f3, Nanog, PCNA and H3K27ac together with H2B, as histones have been shown to compete with these transcription factors for DNA binding^6^) across narrow interphase time windows, where mean PCNA remained stable (Materials and Methods). This analysis revealed that ZGA-associated factors account for a substantial proportion of the residual transcriptional variability independent of CCP (Fig. 4f). The quantified relative contribution of these factors in predicting transcriptional output showed a strong contribution from Pou5f3, followed by PCNA, H3K27ac and Nanog (Fig. 4f’). Finally, we built a comprehensive model that combines the non-linear CCP relationship with additional contributions from ZGA factors as linear predictors (Materials and Methods). These models generalized across held-out embryos (i.e. not used for model training) and more accurately predicted Pol-II-S2p levels compared to models only considering CCP (Fig. 4g-g’ and S6c), even on spatially proximate cells within the same embryo that displayed strikingly different transcriptional activities (Fig. 4h-h’). Together, these results demonstrate that transcriptional heterogeneity during ZGA in early zebrafish embryos emerges from the interplay of within-cycle variations shaped by CCP and across-cycle accumulation of pluripotency factors and chromatin activation marks. By leveraging quantitative, image-based measurements of multiple markers that integrate these processes across scales, our model provides a predictive framework for how individual cells in the zebrafish embryo transition from maternal to zygotic genome activation.

## Discussion

We here extended iterative indirect immunofluorescence imaging (4i)^15^ to confocal *in toto* imaging of early zebrafish embryos (3D-4i), and collected a ten-plex protein dataset in 212 embryos of more than 250 µm in thickness each, resulting in multiplex measurements for ∼10^11^ volumetric pixels. To analyze such terabyte-scale datasets, we built an integrated image analysis pipeline encompassing image pre-processing, nuclei, cell, and embryo segmentation, as well as multiscale feature extraction. Combined with time-course experiments, this offers a comprehensive platform for identifying properties that emerge at different length and time scales as revealed by the presence, abundance, and spatial organization of multiple proteins and protein states in complex 3D systems. By applying this platform to ZGA in early zebrafish embryos, a hallmark developmental transition that is highly heterogeneous between individual cells, we show that it allows the collection of measurements that can train predictive models with high single-cell precision on held-out datasets. This provides a foundation for future machine learning-driven approaches to model cellular decision-making in 3D multicellular systems that is both predictive and mechanistically insightful.

ZGA is orchestrated by multiple molecular processes, including cell cycle progression, chromatin state and the nuclear abundance of pluripotency factors that collectively shape the nuclear and transcriptional landscape of individual cells. Despite much progress in understanding the mechanisms that drive the complex interactions of these factors within individual cells, it has remained unclear what causes the heterogeneous nature of ZGA onset across embryos. By simultaneously measuring cell cycle markers, ZGA-associated factors and transcriptional activity at subcellular resolution across a large number of embryos, we demonstrate that i) cell cycle phase (CCP) can be reliably inferred for every cell within the embryo from fixed samples; ii) CCP is the principal source of variability in zygotic transcription; and that iii) CCP, together with ZGA-associated factors accurately predict transcriptional heterogeneity at single-cell level. The non-linear relationship between CCP and transcriptional activity likely reflects constraints imposed by the DNA replication machinery on chromatin accessibility for transcription, while the additional contribution of ZGA-associated factors underscores their role in modulating transcriptional competence within cell cycle-defined phases. Moreover, these findings suggest that the observed heterogeneity does not arise from imprecise control, but instead emerges from the multimodal causal effects acting at different temporal scales. Future work will be required to determine whether the transient variability in ZGA onset^11^ contributes to longer-lasting patterning establishment after MBT.

Importantly, despite the modest number of proteins and protein states included in our multiplexing panel, we achieved high prediction accuracy with the multivariate set of quantitative features extracted or derived from these stains. This demonstrates that, with careful marker selection and identification of confounding biological processes, such as the cell cycle, highly predictive computational models can be derived even with limited molecular inputs. Nevertheless, further optimization of multiplexed imaging technologies will be essential to expand the number of detectable proteins and post-translational states. Future developments may benefit from complementary preservation strategies, including stronger fixation or embedding methods, as well as improved elution and de-lipidation protocols, to completely abolish background signal accumulation. Ultimately, integrating such higher-plex protein state imaging with multiplexed fluorescent in situ hybridization (FISH)^36^ and single-cell multi-omics data, including chromatin accessibility and transcriptomics^37–39^, will be necessary to enable the construction of comprehensive systems-level models that capture the flow of information across spatial and temporal scales that drive pattern emergence during early development.

## Materials and Methods

### Fish Line Maintenance and Embryo Staging

Zebrafish (Danio rerio) were maintained in the fish facility of Prof. Darren Gilmour. All animal work was carried out by the FELASA guidelines and standards of the University of Zurich. Embryos were raised at 28 °C in E3 medium until reaching the corresponding developmental stage. During sample collection, initial embryo staging was performed according to time post fertilization^40^. The experiments were carried out on Pou5f3-3xflag transgenic line, which was generated by Dr. Edlyn Wu in the laboratory of Prof. Nadine Vastenhouw at the University of Lausanne according to Swiss regulations (canton Vaud, license number VD-H28).

### Generation of *pou5f3:3xflag* zebrafish line

The 3xflag knock-in *pou5f3* line was generated using CRISPR/Cpf1^41^. Guide RNA (gRNA) was designed using CHOPCHOP tool^42^ and targets sequence close to the C-terminal end of *pou5f3* gene 18 bp upstream of the stop codon. The sequence of gRNA is as follows (with PAM sequence underlined): TTTGGTGGGTCATCTCACCAGCTAACTT. The sequence of DNA repair template (donor ssODN) is as follows (with mutated PAM sequence underlined, 3xFLAG sequence in brackets, and stop codon in bold): TGCCATCCCTCCACCGACCAGATGTCTTCAAAAACGGCTTGCACCCTGG<u>TCTA</u>GTGGGTCATCTCACCAGC[G ACTACAAAGACCATGACGGTGATTATAAAGATCATGATATCGATTACAAGGATGACGATGACAAG]**TAA**CTTGCT CCAGGTGCTCACTAAAGTTTATAGAAA (173 nts, synthesized by IDT).

RNP mix was first prepared as follows: 1 μl EnGen LbaCas12a (Cpf1) protein (M0653T, NEB); 0.353 μl gRNA (at 100 μM stock), 5 μl RNP buffer (50 mM KCl, 2.5 mM MgCl_2_, 5 mM Tris-HCl pH 8), and 1.65 μl nuclease-free water. RNP mix was incubated at 37°C for 10 mins and then kept at room temperature before use. To prepare Cpf1 injection mix, 4 μl RNP mix was used and 0.5 μl 400ng/μl ssODN and 0.01% phenol red were added. A final amount of 150 pg gRNA, 40 pg ssODN, and 20 fmol Cpf1 was injected into the one-cell stage of zebrafish embryos. Injected embryos were grown for three months at 28°C before screening for mutant founders. Founders were outcrossed to wild-type (TL/AB) fish. To obtain homozygous fish, outcrossed fish originating from two founders were in-crossed. Homozygous fish were screened by finclipping. Homozygous *pou5f3-3xflag* embryos were produced by in-crossing identified homozygous fish.

### Sample Preparation

*pou5f3-3xflag e*mbryos were collected in 15-minutes intervals between 2 and 4 hours post-fertilization (hpf) and fixed in fresh and pre-cooled 4% paraformaldehyde (PFA; Lucerna Chem) in 1X PBS. Tubes were kept on ice during sample collection and were then transferred to a rotary shaker at 4°C for overnight fixation (up to 18 hours). Fixation was halted by thoroughly washing the embryos five times with 1X PBS, followed by quenching with 100 mM ammonium chloride (Sigma-Aldrich) for 30 minutes. Embryos were then dechorionated in a glass Petri dish using watchmaker forceps under a stereo microscope and permeabilized by gradually increasing the methanol concentration in 25% increments up to 100% (Sigma-Aldrich), with 5-minute intervals between each step. The embryos were then incubated in 100% methanol at -20°C for 2 days. Successful storage was tested for up to 3 months. On the experiment day, samples were re-hydrated back to PBS-T (PBS+0.1% Tween 20; Sigma-Aldrich) in 25% steps. Consequently, yolk was removed by agitating the samples in a glass centrifuge tube using a Pasteur pipette in at least six rounds of washing with PBS-T.

### Mounting on 96-well Plates

Plate preparation was done on the day of mounting. The wells of a polystyrene, flat-bottom 96-well plate (Greiner) were coated as follows to facilitate sample adherence. The coating solution consisted of a 1:10 dilution of human fibronectin (Sigma-Aldrich) in PBS mixed with a 1:2 dilution of Poly-D-Lysine (Thermo Fisher) in PBS. Each well received 50 μl of the solution and was incubated for two hours at room temperature. After incubation, the liquid was aspirated, and the plate was left to dry for an additional two hours. Wells were filled with 300 μl PBS, then the embryos were carefully transferred into the wells using a glass Pasteur pipette with a Pipet Pump II (Sigma-Aldrich) to enable gentle handling. Allowing the embryos to settle naturally within the liquid column of the filled well ensured their robust orientation with the animal pole facing the bottom of the plate. Up to 9 embryos were mounted in a single well, to maintain proper spacing between them for imaging. Embryos from different time points were pooled and randomly distributed within wells to alleviate potential sources of technical bias between different developmental timepoints. After mounting, the plate was incubated for two hours before centrifugation at 200 rcf for five minutes on a plate centrifuge. To enhance adherence, the plate was placed on a UV transilluminator (Lab Gene Instruments) for one hour at 365 nm with high intensity for crosslinking. It was then stored at 4°C overnight before proceeding with the next steps. For additional crosslinking, an imaging session was performed on the Cell Voyager 8000 microscope, acquiring a 100 μm stack with a z-spacing of 4 μm using the 488 nm laser at 100% laser power and 100 ms exposure.

### Whole-mount Multiplexed Immunofluorescence

The protocol for performing 3D-4i on zebrafish embryos was devised based on standard immunofluorescence protocol for zebrafish embryos^43^, the original 4i method^15^ and whole-mount volumetric imaging^44^. Embryos were prepared and mounted in a 96-well plate as described before. Each cycle of the multiplexing experiment consisted of the following steps, all conducted at room temperature:

1. Antigen Retrieval/Antibody Elution
2. Blocking
3. Primary Antibody Incubation
4. Secondary Antibody Incubation
5. Optical Clearing by Refractive Index (RI) Matching
6. Imaging
7. RI Matching Buffer Washout

All pipetting steps were executed using Bravo automated liquid handling platform (Agilent) with protocols described below. A residual volume of 70 μl was always maintained in each well to avoid sample disturbance when aspirating/dispensing liquid.

#### 1. Antigen Retrieval/Antibody Elution

The 3D-4i elution buffer (similar to 4i elution buffer: 0.5 M L-Glycine (Sigma-Aldrich), 1.2 M Urea (Sigma-Aldrich), 3 M Guanidinium chloride (Sigma-Aldrich)) was prepared in advance, with Tris(2-carboxyethyl)phosphine hydrochloride (TCEP; Sigma-Aldrich, 0.07 M) added just before use. The pH was then adjusted to 3 using 32% HCl (Sigma-Aldrich). The buffer was introduced via multiple volume replacement and mixing steps. After incubating for 1 hour, wells were thoroughly washed 12 times with PBS (total duration of 30 min), as such pH returned to approximately 7.

#### 2. Blocking

The 3D-4i blocking buffer (PBS with 1% BSA and 0.2 M Ammonium chloride; Sigma-Aldrich) was prepared at 2X concentration in advance. Before use, Maleimide (0.3 M; Sigma-Aldrich) was added. The blocking buffer was mixed with the equal volume of residual PBS in the wells and embryos were incubated in the blocking buffer for 1 hour, followed by extensive washing with PBS.

#### 3. Primary Antibody Staining

##### 3.1 Indirect Immunofluorescence

Primary antibodies were prepared at 2X concentration in a 2X 3D-4i blocking buffer (without Maleimide) and mixed with an equal volume of residual PBS in each well. Samples were incubated in this solution overnight at room temperature on a gentle see-saw rocking device.

##### 3.2 Direct Immunofluorescence

Antibodies against high-abundance targets showed strong staining bias from periphery to the center of the sample^45^. To alleviate this issue, these primary antibodies were directly labelled with fluorescently tagged Fab fragments. The ZenonTM IgG labelling kits (Invitrogen) were used as indicated by the manufacturer’s protocol. In short, 1 μg of primary antibody was prepared and incubated with 5 μl of Zenon Fab fragments for 5 minutes. Surplus unbound Fab fragments were blocked using 5 μl of corresponding species blocking buffer provided in the kit, incubating for another 5 minutes. Labelled primary antibodies were prepared fresh for each staining and incubated overnight with embryos at room temperature. No additional secondary step was performed, except for DAPI staining prior to imaging. List and dilution factors for each primary antibody and the cycle in which they were stained, are listed in Table1.1.

**Table1.1:** List of primary and their corresponding secondary antibodies used in the multiplexing experiment, their dilutions and staining cycles.

| Primary | Host | Dilution | Stain Type | Secondary | Secondary Dilution | Staining Cycle | Catalogue Number | Vendor |
| --- | --- | --- | --- | --- | --- | --- | --- | --- |
| Pol-II-S2p | Rat | 1:250 | Indirect | Donkey anti- Rat 488 | 1:200 | 1 | 04-1571 | Sigma-Aldrich |
| Anti-FLAG | Mouse | 1:500 | Indirect | Donkey anti- Mouse 568 | 1:200 | 1 | F1804 | Sigma-Aldrich |
| PCNA | Rabbit | 1:250 | Indirect | Donkey anti- Rabbit 647 | 1:200 | 1 | 13110 | Cell Signaling Technology |
| H3K27ac | Rabbit | 1:500 | Direct | Zenon Rabbit 488 | 1:100 | 2 | ab4729 | Abcam |
| β-Catenin | Mouse | 1:250 | Indirect | Donkey anti- Mouse 568 | 1:400 | 2 | 2677 | Cell Signaling Technology |
| pH3 | Rabbit | 1:150 | Direct | Zenon Rabbit 647 | 1:150 | 2 | 9701 | Cell Signaling Technology |
| H2B | Rabbit | 1:100 | Direct | Zenon Rabbit 488 | 1:250 | 3 | ab134211 | Abcam |
| ALYREF | Mouse | 1:250 | Direct | Zenon Mouse 555 | 1:100 | 3 | sc-32311 | Santa Cruz Biotechnology |
| Pol-II-S5p | Rat | 1:250 | Indirect | Donkey anti- Rat 647 | 1:200 | 3 | 04-1572 | Sigma-Aldrich |
| Nanog | Rabbit | 1:280 | Direct | Zenon Rabbit 488 | 1:28 | 4 | GTX132491 | GeneTex |

We note that Anti-FLAG antibody staining in *pou5f3-3xflag* embryos serves as a readout for Pou5f3 protein abundance.

##### 4. Secondary Antibody Staining

Excess primary antibodies were removed by extensive washing with PBS. Secondary antibodies were diluted at 2X concentration in 3D-4i blocking buffer (without Maleimide) and supplemented with DAPI (final dilution: 1:40; Invitrogen). The 2X antibody and DAPI mixture was combined with an equal volume of the residual PBS in each well and incubated on samples for 4 hours under gentle agitation in the dark. Details for each secondary antibody and Fab labeling kit are listed in Tables 1.1 and 1.2.

**Table1.2:**
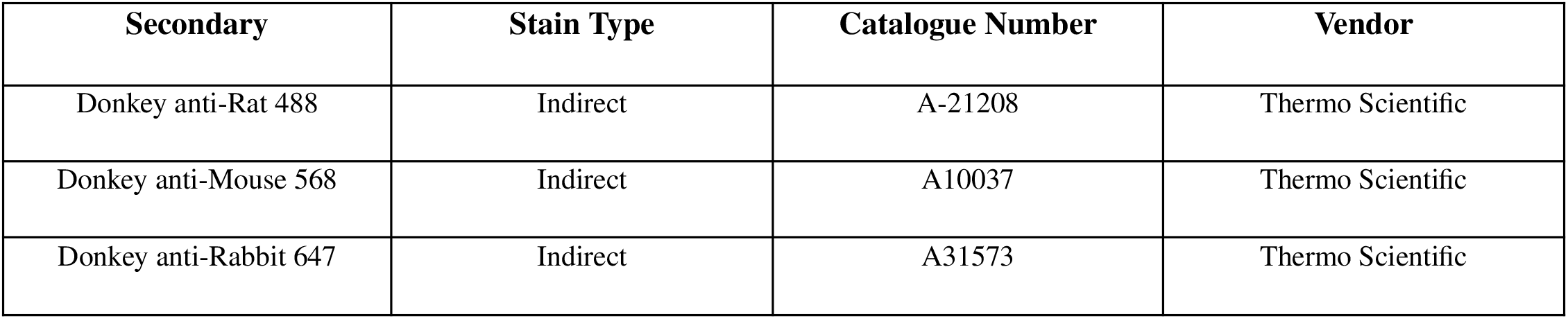
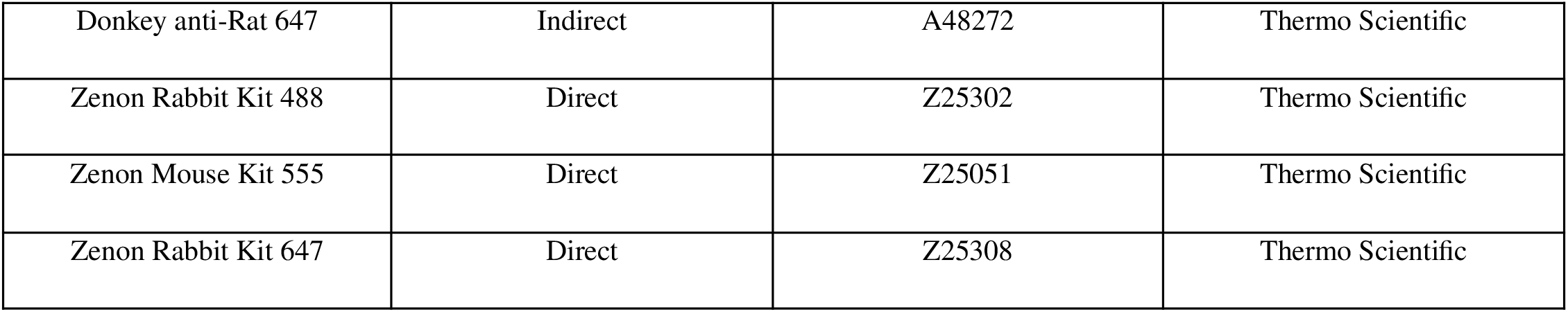
List of secondary antibodies and Fab fragment labeling kits used in the multiplexing experiment.

#### 5. Optical Clearing by RI Matching

Excess secondary antibodies and dyes were removed by extensive washing with PBS. Optical clearing solution for refractive index matching was prepared as advised by Chung lab protocol: https://docs.abcam.com/pdf/protocols/clarity-protocol.pdf. N-acetyl-cysteine (Sigma-Aldrich, 0.7 M) was added to this buffer prior to imaging to reduce oxidative damage caused by light exposure during imaging. Lastly, DAPI was added to the imaging buffer in the same concentration used for staining, to prevent its gradual diffusion out of the sample during long imaging time. Embryos were kept in the imaging buffer for at least 1 hour prior to imaging to ensure buffer equilibration.

#### 6. Imaging

Imaging was performed using a Yokogawa CV8000 spinning disk (CSU-W1) inverted confocal microscope equipped with a 50 μm pinhole disk and a Hamamatsu ORCA Flash 4.0 camera. A 20x/1.0 NA water immersion objective was utilized to capture high-resolution images, with each field of view containing a single embryo. Embryos were detected and selected using a custom Python script. A z-stack of 271 μm was acquired with 1 μm spacing between optical sections. Laser power and exposure time were adjusted individually for each stain to optimize signal quality. Imaging was conducted overnight.

#### 7. RI Matching Buffer Washout

The imaging buffer was washed out thoroughly using PBS after each cycle of imaging.

### Re-staining Controls

As a technical control, markers from the first three staining cycles were re-stained in cycle four, each using three replicate wells. Signal correlations between the original stains and their re-stainings were assessed at the mean nuclear level, and at the single-pixel scale using full-resolution or two times down-sampled images (Fig. 1f-g).

### HDF5 format

An important step in standardizing image analysis workflows is the on-disk specification of data. To streamline this process, we developed a method for storing image data using a custom container based on HDF5. This format is designed to store image pyramids (multiple down-sampled resolutions of raster data) and label images. Here, each microscopy site is written to a HDF5 file, containing groups for each channel and its corresponding resolution levels. Metadata are stored as dataset-level attributes. In the past years, OME-Zarr has emerged as a standard format across the open-source bioimage community^46^ and is thus recommended for new projects.

### Image Registration of Acquisition Cycles

During each cycle of immunofluorescence imaging, sample shrinkage and slight displacements occur due to exposure to elution and imaging buffers and variations in microscope stage initializations, respectively. This necessitates an unsupervised approach to register all acquisitions within a multiplexing experiment to a reference acquisition cycle. To address this, we stained the samples with DAPI in each cycle to label DNA, which serves as a reference channel for registration across cycles. We then utilized the elastix library^21,22^, which offers a comprehensive suite of tools for developing custom image registration workflows. Due to embryo shape deformations and stage shifts across cycles, we applied three sequential registration steps, each introducing more degrees of freedom for alignment: starting with rigid (translations and rotations), progressing to affine (rigid, scale, and shearing), and finally applying b-spline (elastic, non-linear transformation) registrations (Fig. S1a). The second acquisition cycle was selected as the reference cycle for registration, as it contained the cell segmentation marker (β-catenin). Although these transformations significantly improved the alignment of embryos at single-cell resolution across cycles (Fig. S1), some nuclei, particularly those at the borders of the imaged volume, remained misaligned. To assess alignment performance quantitatively, we measured Pearson’s correlation coefficient of nuclear DAPI between cycles. This coefficient was above 0.9 for most cells, and cells with a correlation below this threshold were excluded from downstream analysis (Fig. S1b).

### Object Segmentation

Embryos were segmented using β-catenin and DAPI staining and by training a pixel classifier in Ilastik^23^. Nuclei segmentation was performed using the pre-trained Cellpose nuclei model with the expected nuclei diameter set to 25 pixels^24^ (=16.25 µm). To ensure consistent nuclei segmentation across the embryo, the DAPI signal was normalized along the z-axis by applying an exponential rescaling correction with a factor of -0.0045, which improved signal intensity at higher z-planes. This correction was applied solely to enhance segmentation performance; raw DAPI intensities were left unmodified for downstream analyses. For cell segmentation, we implemented a solidity-constrained seeded watershed approach, with nuclei as seeds and embryo mask as foreground, using the scikit-image implementation^25^. Membrane labels were generated by selecting the single-pixel outline of the cell labels, and cytoplasm segmentation was obtained by subtracting both the nuclear and membrane labels from the cell masks. Finally, for subnuclear detection of active transcription sites a second pixel classifier was trained using Pol-II-S2p staining in Ilastik^23^. Segmentation steps were performed on two times down-sampled images, as this resulted in a more isotropic x, y, z voxel spacing compared to the full-resolution images, thereby improving segmentation performance.

### Features Aggregation and Extraction

By aggregating features across multiple levels, from individual voxel measurements to objects and hierarchical interactions, we gain a comprehensive understanding of complex biological systems. This approach allows us to study not only individual cells but also their broader tissue context, providing powerful insights into spatially resolved biological data. The structured organization of features into aggregating tables ensures efficient querying, aggregation, and integration into machine learning workflows, making this process scalable for large datasets. This approach is broken down into the following layers:

1. Voxel-level features: Each voxel carries information about its location, extent, fluorescence intensity, and its place in the hierarchical object structure (e.g., nucleus, cell, embryo). These voxel properties serve as the foundation for raster features, such as volume, intensity, and shape (e.g., roundness).
2. Raster features form the most fundamental class of features and are the basis for all downstream measurements. These features include both label-based (nucleus, cell, etc) and intensity-based measurements
  A. Label features: These are computed from label images and include label ID, bounding box, centroid, and morphological properties such as roundness, volume, and Feret diameter^26^.
  B. Intensity features: These features describe the intensity distributions within label regions using descriptive statistics (mean, median, kurtosis, etc.)^26^.
  C. Colocalization features: These features measure the relationship between staining channels within label regions, quantifying the correlation or mutual information between different fluorescence markers and are useful for studying protein interactions and validating registration quality.
  D. Distance features: These features calculate distances between label and reference objects (e.g., from a nucleus to its parent cell). They include centroid distances, maximum, minimum, and median distances based on Euclidean distance transforms.
3. Population features focus on spatial interactions between objects represented as a graph, where nodes are objects and edges depict spatial or hierarchical relationships. We construct four distinct types of graphs:
  A. Radius neighborhood: Objects within a given radius are considered neighbors.
  B. K-nearest neighbors (KNN): Each object is connected to its k closest neighbors, regardless of their distance, ensuring a fixed degree for all nodes.
  C. Delaunay triangulation: Neighbors are defined by applying Delaunay triangulation on objects, partitioning the convex hull into tetrahedra.
  D. Touch neighborhoods: Adjacency is defined by shared surface area between objects.
4. Hierarchy feature aggregations take place across different levels of the object hierarchy. For instance, by aggregating cells to their parenting embryo, average cell cycle phase or number of cells in an embryo can be measured. This level provides a global view of embryos/tissues.
5. Feature embeddings include combining multiple feature columns into higher-level properties. These embeddings offer compact representations of complex data and can be either categorical or continuous.

### Intensity Bias Corrections

To estimate flatfield corrections, the following fluorescent dyes were diluted in DMSO, mixed and added to a Greiner 96-well plate: 7-diethylamino 4-methyl coumarin (Sigma-Aldrich, 0.5 mg/ml), Fluorescein (Sigma-Aldrich, 1 mg/ml), Rose Bengal (Sigma-Aldrich, 1 mg/ml) and acid blue 9 (TCI Chemicals, 10 mg/ml). Multiple imaging stacks were acquired for each wavelength, following the same acquisition protocol as described earlier. Flat fields for each channel were then calculated using the BaSiC Fiji plugin^28^. Dark fields were recorded by averaging 100 images with the laser turned off. Dark and flat field corrections were applied irreversibly during the conversion from raw TIFF images to HDF5 format. The corrected images were clipped at 0 to prevent integer overflow before being cast to uint16. Acquisition time (t-decay) and z intensity (z-decay) bias models were estimated based on the mean intensity features extracted after nuclei segmentation. Both t-decay and z-decay models were estimated separately for each stain, by fitting a single or double exponential curve to the data. Bias corrections were applied to the imaged volumes on the fly prior to final feature extraction. Background signal was estimated by measuring the mean signal intensity in the cytoplasm. This was then subtracted from the mean nuclear signal to obtain mean background-subtracted values. The sum nuclear intensity was subsequently calculated by multiplying the background-subtracted nuclear mean to nuclear volume.

### Napari Feature Classifier

We developed a napari feature classifier plugin for annotation and interactive classification of labeled objects: https://napari-hub.org/plugins/napari-feature-classifier.html. The plugin supports loading measurements and attaching them to a label image and its corresponding multi-channel images in napari. Users define classes and interactively assign training samples, after which a random forest classifier is trained within the interface. Predicted object classes are displayed in real time, enabling iterative refinement of the training set by adding examples in regions where classification performance is sub-optimal. We used this plugin to classify segmented nuclei to debris, enveloping layer (EVL), yolk syncytial layer (YSL), or deep cell labels.

### Cell Cycle Phase (CCP) Inference

The following steps were involved in generating the CCP embedding and hyperparameter tuning:

1. Feature pre-processing:
  A. Outlier removal: Removing label objects with fluorescence artifacts in the image channels or mis-segmented nuclei.
  B. Stratification: Stratifying the data into division cycle bins based on estimated embryo age, defined as log2 of nuclei count. Embryos from cycles 7 to 9 were pooled due to fewer cells at these stages.
  C. Sub-sampling: Ensuring an equal number of cells per stratum.
2. Pre-embedding:
  A. Feature selection: From the complete feature panel, we selected multiple nuclear morphology features (nuclear volume, roundness and flatness and nuclear/cytoplasmic ratio) as well as intensity features of cell cycle markers DAPI sum (DNA content) and mean PCNA (ccp-features in table 2). pH3 was excluded from the features in the final model due to background accumulation.
  B. Feature scaling^47^:
    a. Standard scaler: Rescaling to 0 mean and 1 standard deviation.
    b. Robust scaler: Rescaling to 0 median and 1 quantile range.
    c. Power transformer: A monotonic non-linear transformation toward a Gaussian distribution with 0 mean and 1 standard deviation using the ’Yeo-Johnson’ method.
  C. Pre-embedding:
    a. An embedding method was selected from scikit-learn compatible dimensionality reduction algorithms (e.g., PCA^48^, UMAP^33^, or Palantir^49^).
    b. Embedding dimensionality was specified, with 2-4 dimensions evaluated.
    c. Stratification was optional, see “Cell Cycle Phase Scoring Pipeline and Hyperparameter Tuning”.
3. CCP inference:
  A. Fit a principal circle to the pre-embedded nuclei of each stratum^50^.
  B. CCP embedding: Compute this by fitting a smooth b-spline through the principal circle nodes.
  C. CCP inference: Approximate the CCP for each nucleus using:
    a. Distance-based inference, where CCP is defined as the position of the closest point on the circle using Euclidean distance.
    b. Angle-based inference (only for 2D pre-embedding), where CCP is defined as the angle to the center of mass of the loop.
  D. Alignment:
    a. Detect peaks and infer their identity (S to M phases) and order using cell cycle markers.
    b. Use peak values in the 2^nd^ derivative of mean PCNA intensity to identify the start and end of the S phase.
  E. Normalization:
    a. To achieve a uniform density distribution for each stratum, we applied the QuantileTransformer from Scikit-learn.

**Table2:** Feature sets used in regression modeling. Linear predictors included CCP, cell cycle features and ZGA factors. Non-linear CCP predictors included B-spline, asymptotic, and power regressions. Model names indicate included feature sets, e.g., *zga_ccp-asympt* combines four ZGA predictors with asymptotic CCP regression (7 parameters in total).

| Component | Features | Description |
| --- | --- | --- |
| **Regression target** |  |  |
|  | Pol2.S2p | Pol-II-S2p (transcription elongation) |
| **Linear predictors** |  |  |
| ccp | CCP | Interphase CCP |
| ccp-features<br>(ccp feats) | PCNA, DAPI, nuclei volume,<br>nucleus/cytoplasmic volume ratio,<br>nuclei roundness, nuclei flatness | A subset of features that were used for cell cycle phase inference. |
| zga | Pou5f3, Nanog, H3K27ac, H2B | ZGA-associated factors |
| **Non-linear predictors** |  |  |
| ccp-asympt | CCP (a,b,c), equation 4 | Asymptotic regression on CCP |
| ccp-power | CCP (a, b), equation 5 | Power curve regression on CCP |
| ccp-spline | Bs (CCP, 3, fit_intercept=False) | B-spline encoded interphase CCP (3 knots without intercept) |

### Cell Cycle Phase Scoring Pipeline and Hyperparameter Tuning

To find the optimal CCP inference, we included as many hyperparameters as possible in the above-mentioned pipeline and evaluated these models, by manually classifying 634 cells across different time points into 6 cell cycle classes (anaphase, telophase, early S phase, S phase, prophase, and metaphase of mitosis, as shown in Fig. 3b-b’). This manual cell cycle annotation was performed using DAPI, PCNA and pH3 channels, nuclear morphologies and embryonic context (e.g. mitotic cells in the vicinity) as visual metrics. We assessed the rank ordering of the models by partitioning the model-inferred position of the annotated cells within the embedding into six classes and counting the number of pairwise permutations required to achieve the ground-truth annotated classes. A challenge arose due to the circular nature of the embedding, where the numerical values of 0 and 2π correspond to the same point, meaning metaphase and anaphase are adjacent. To account for this, instead of the usual 5 cuts to partition a one-dimensional feature into six regions, we required six cuts for this circular feature. Each model was evaluated for all strata (cycles 7-9, 10, 11 and 12), and its worst score across these cycles was recorded to ensure the selection of a model that performs reasonably well across all embryonic stages. Each run involved defining a parameter grid and evaluating the resulting models based on the scoring procedure. The key trends that guided the selection of the final model were:

1. **Sub-sampling** was the most significant contributor to model performance, particularly at later cycles that are heavily skewed toward interphase. Such class imbalance hinders correct CCP inference across all phases.
2. **Non-linear feature scaling** outperformed both robust and standard scaling approaches.
3. **Non-linear pre-embeddings** such as UMAP and Palantir outperformed linear ones e.g. PCA.
4. **Pre-embedding stratification** per division cycle slightly outperformed a single pre-embedding.
5. **Separate principal circle fitting** outperformed a single fit for all strata.

Since subsampling emerged as a major determinant of model performance, we repeated the procedure by using the initially selected CCP model to guide a new round of data subsampling. The model was then retrained on this resampled dataset, which yielded qualitatively improved results.

We note that although the global structure of the CCP trajectory resembles cell cycle progression, as indicated by the gradual doubling of total DNA content during interphase (DAPI sum, Fig. 4a), the local positioning of cells within the CCP may contain errors. For example, some mitotic nuclei resembling the metaphase-to-anaphase transition are positioned at prophase according to CCP inference, likely due to their morphological similarity to prophase nuclei. Such errors should not influence downstream prediction analyses, as mitotic cells were excluded.

### Embryo Staging

Since embryos of different timepoints were mixed in the same well to minimize technical variability, they were first re-staged based on nuclei counts during data analysis steps (Fig. 2i). However, this method provides only cycle-level resolution, and missing nuclei can be misinterpreted as differences in developmental age. To overcome this limitation, following nuclei count-based staging, embryos were further ordered according to their circular mean CCP, thereby refining developmental staging (CCP-adjusted staging, Fig. S4a).

### Cell Cycle Phase Circular Statistics

CCP is a circular variable ranging from 0 to 2π, with the endpoints representing the same phase of the cell cycle. To account for this periodicity, we used Scipy’s circular statistics^51^: the circular mean Θ (scipy.stats.circmean) to compute the average CCP per embryo and circular synchrony R as R= 1-scipy.stats.circvar, measuring the reduction in CCP variance across cells within each embryo (Fig. S4 and 3e). To estimate local CCP synchrony (Fig. 3f), “R” was calculated by aggregating cells in radius-defined neighborhoods ranging from 0 to 500 μm.

### Modeling Global Cell Cycle Synchrony

We modeled R as a function of CCP-adjusted staging (s) using a four-parameter Hill equation:

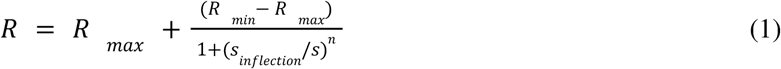

where *R* _*max*_, *R* _*min*_ are the asymptotic maximum and minimum synchrony, *s*_*inflection*_ is the stage at half-maximal synchrony and *n* is the Hill coefficient. Parameters were estimated by non-linear least squares using SciPy’s scipy.optimize.curve_fit. Pointwise confidence intervals for the mean Hill curve were obtained via a residual bootstrap, where after the initial fit the residuals (observed R minus fitted R) were resampled and added to the fitted values to create synthetic responses (200 datasets). The Hill model was refit for each bootstrap sample and the resulting mean and 95% confidence intervals were reported.

### Cell Cycle Phase Gradient Estimation and Pseudowave Reconstruction

CCP gradients were computed using a finite difference method designed for irregularly spaced data^52^. The method was adapted to operate on 3D point clouds and modified to incorporate circular differences, enabling application to CCP. The resulting gradient field was integrated with PyVista to generate streamlines for visualizing mitotic pseudowaves (Fig. 3g).

### Modeling of Transcriptional Activity within Division Cycles

We modeled transcriptional activity during S phase using natural log-transformed mean nuclear Pol-II-S2p intensity, as a proxy for transcriptional output^3,19^ by performing multilinear regression. The beginning and end of the S phase was defined using peak values in the 2^nd^ derivative of mean PCNA intensity. Regression models were formulated in a Bayesian framework (using PyMC^53^ and Bambi^54^) with posterior inference obtained by Markov Chain Monte Carlo sampling, and model diagnostics performed using ArviZ library^55^. The response variable, y, was assumed to be normally distributed:

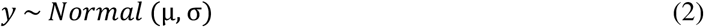

with expected value μ and variance σ. We tested a range of model specifications for their expected values. Linear models included predictors (*X*_*limear*_ ) such as CCP and ZGA factors (zga; including Pou5f3, Nanog, H3K27ac, H2B). Additionally, we included features used for CCP inference as linear predictors in our modeling approach (ccp-feats: PCNA and DAPI intensities, nuclear volume and shape descriptors, see Table 2), as they may capture additional information beyond that represented by CCP. These linear models are depicted as:

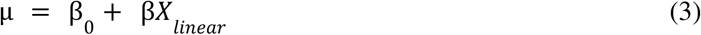

with β_0_ and β assuming to follow normal distributions.

As the relationship between CCP and Pol-II-S2p was found to be non-linear (Fig. 4b), we additionally implemented asymptotic, power (equations 4 and 5) and b-spline (3 knots without intercept) regression models of *X*_*CCP*_ :

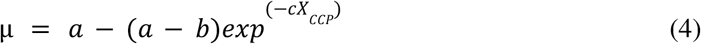

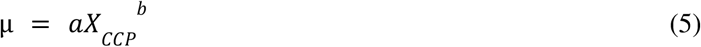

Finally, these non-linear CCP formulations were also combined with linear predictors to capture all features that vary non/linearly in explaining transcriptional activity, as shown for asymptotic and power models below:

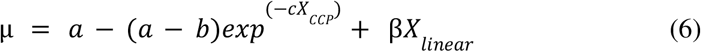

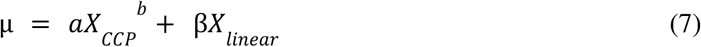

with coefficients a, b and c assuming to follow normal distributions.

Model names indicate included feature sets. For example, the model *zga_ccp_ccp-power* combined linear predictors (ZGA factors and CCP) with a non-linear power regression on CCP. For the asymptotic and power regression models, we tried additional models by clipping the data at 100 gray values as detection accuracy drops at these low signal-to-noise ratios (termed ccp-cens-asympt, ccp-cens-power).

Intensity features used for prediction were first natural log-transformed and, together with morphology features, z-scored to enable direct comparison across features. Models were trained on up to 500,000 cells per division cycle (cycles 8-12). Predictive performance was assessed using explained variance (R^2^) and expected log pointwise predictive density (ELPD).

The best performing model, zga_ccp-feats_ccp-asympt, was trained for each division cycle separately and evaluated on exemplary embryos excluded from training steps in that cycle (Fig. 4g-g’).

### Modeling of Transcriptional Activity across Division Cycles

To model transcriptional activity across division cycles, we divided S phase into six equally sized strata during which the S phase marker PCNA remained unchanged. Within each stratum, with CCP held constant, we trained multilinear regression models to predict transcriptional output (natural log-transformed mean nuclear Pol-II-S2p intensity) from z-scored and natural log-transformed ZGA factor levels, enabling direct comparison of regression coefficients. Model performance was evaluated using 5-fold cross-validation, and predictive accuracy was summarized by R^2^ calculated from the combined S phase data. Regression coefficients for each stain were estimated separately within each stratum and their distributions are shown in Fig. 4f’.

## Data and Code Availability

All codes and data generated in this study will be publicly available upon manuscript publication.

## Figures

**Fig. S1:**
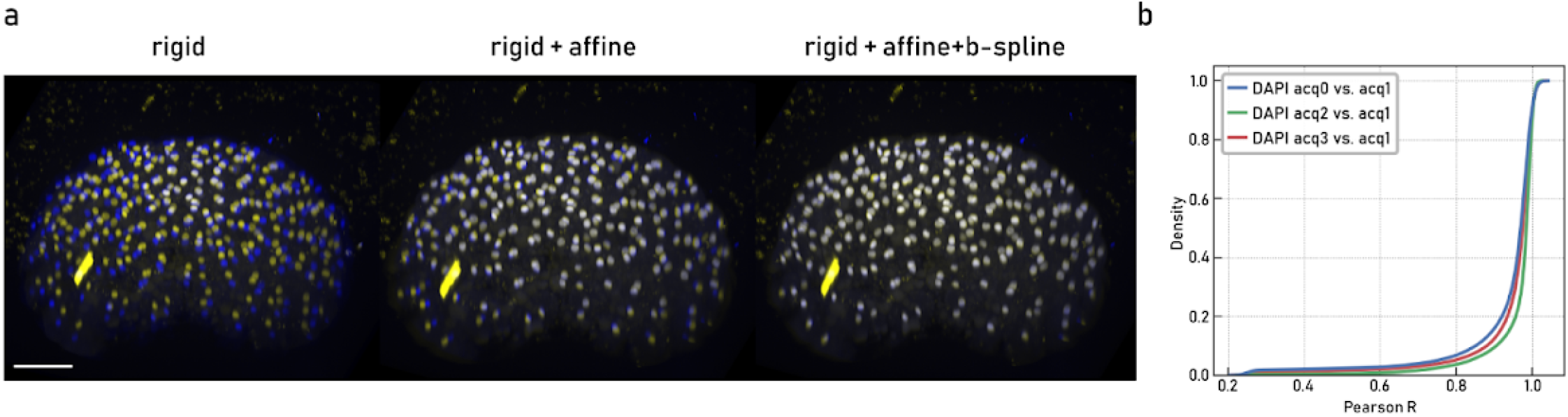
Acquisition cycle registration. a. DAPI stainings of a representative embryo imaged across two acquisition cycles (blue is the reference cycle and yellow the aligned cycle) after rigid (left), rigid and affine (middle) or rigid, affine and b-spline registration (right) steps. b. Cumulative distribution of nuclear DAPI correlations from acquisition (acq) cycles 0, 2 or 3 with the reference cycle 1 across all embryos after registration, used as single-cell estimates of registration fidelity.

**Fig. S2:**
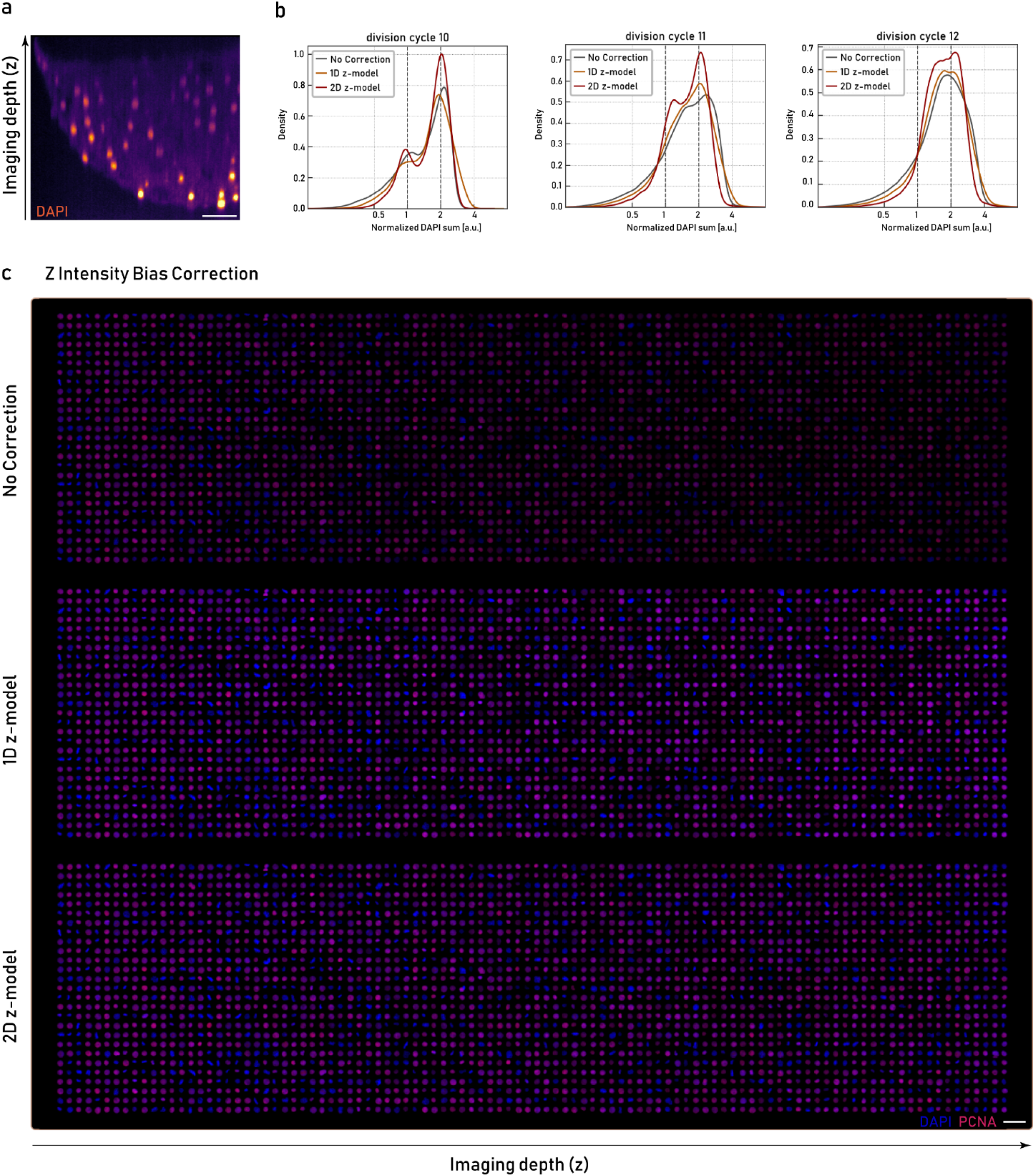
Z intensity bias correction. a. A representative embryo stained with DAPI showing intensity decay across the imaging depth/z axis (animal pole is at the coverslip). Scale bar 50 µm. b. The density plot on comparison of 1D and 2D z-decay models with uncorrected data for summed DAPI intensity of embryos in cycles 10 to 12. Data normalized to the first peak. c. Maximum intensity projections of representative nuclei stained with DAPI (blue) and PCNA (red). Top: uncorrected data, middle: after 1D z-decay correction and bottom: after 2D z-decay correction. Nuclei are sorted by z-coordinate (left to right corresponds to increasing imaging depth (z)). Scale bar 50 µm.

**Fig. S3:**
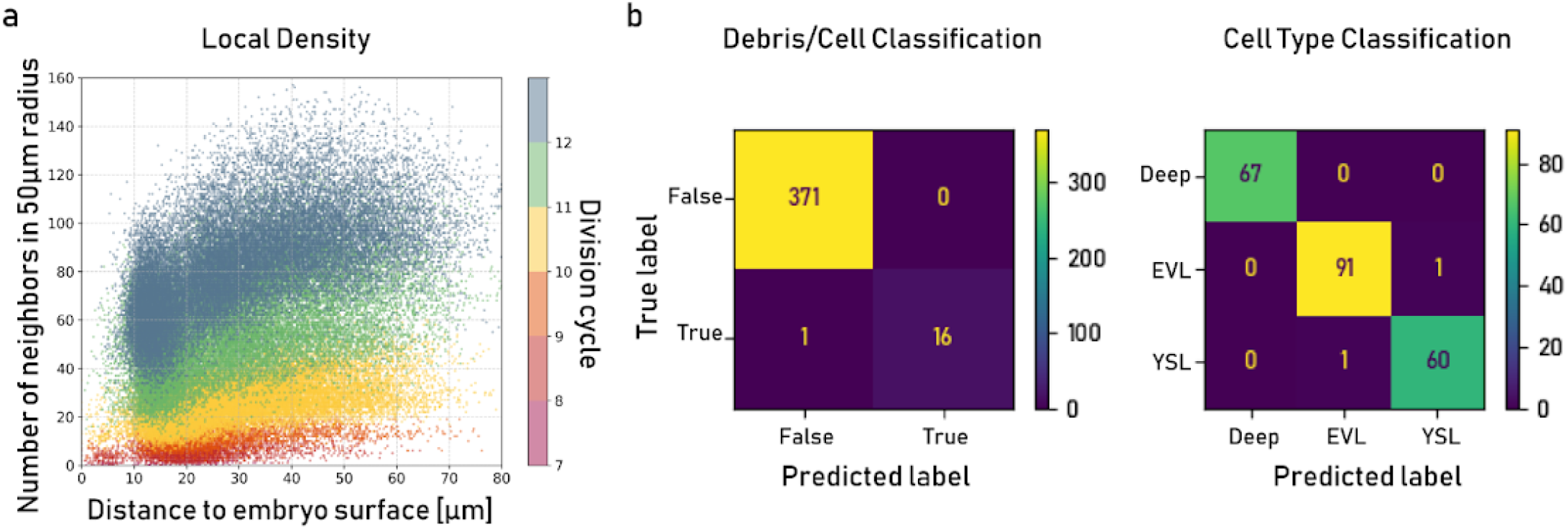
Local density and label object classification. a. Number of neighbors in a 50 µm radius for each cell plotted against its distance to the embryo surface. Results are color-coded per division cycle. Maximum 30,000 cells for each cycle are shown. b. Confusion matrices for the two napari feature classifiers, debris vs. cells (left) and deep cells vs. EVL and YSL (right), evaluated on the test set.

**Fig. S4:**
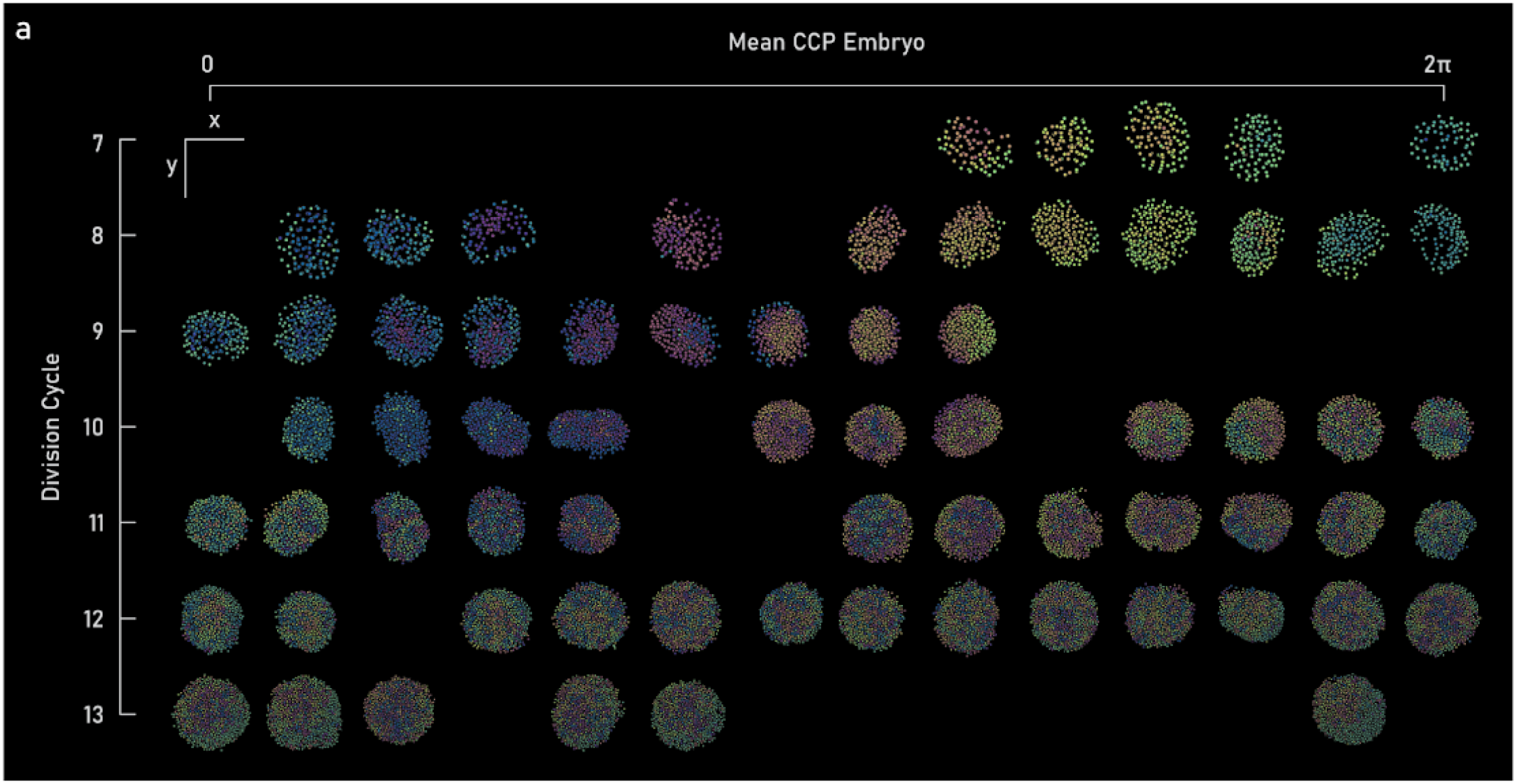
Refined embryo staging. a. Top view (along the z-axis) of a subset of embryos across the full developmental time course studied here, with nuclei colored by inferred CCP. Embryos are ordered by division cycle (top to bottom) and CCP circular mean (left to right). Each grid point corresponds to an average CCP bin and a division cycle; empty bins indicate no matching embryo. Scale bars: 550 μm (x and y).

**Fig. S5:**
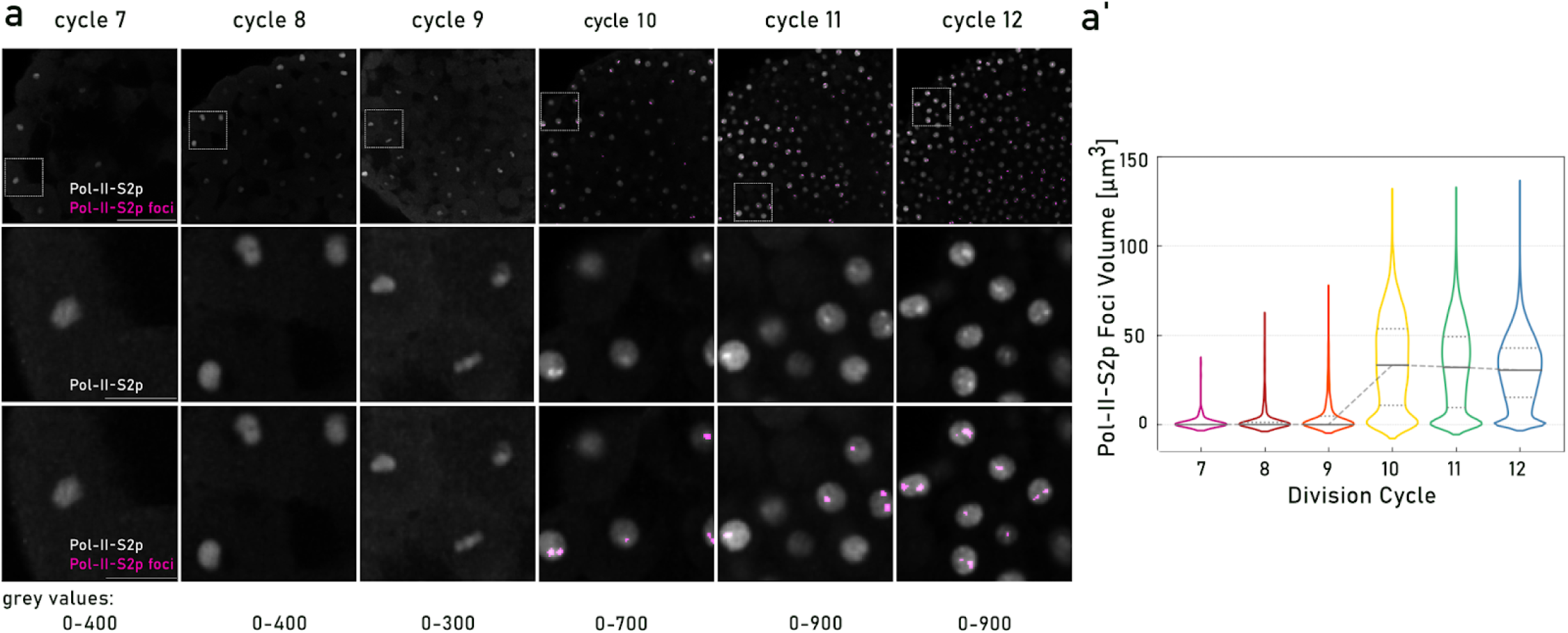
Active transcription sites measurements. a. 2D sections of representative embryos stained with Pol-II-S2p antibody across division cycles 7 to 12, overlaid with active transcription foci labels detected by Ilastik pixel classifier (top and bottom rows). The last two rows demarcate the zoom-in views of the ROIs shown in top panels. Scale bars in overview and zoom-in views are 100 and 25 µm, respectively. a’. Active transcription foci volume measurements for all deep-cell nuclei in S phase across division cycles 7 to 12. The dashed line connects the median values for each division cycle.

**Fig. S6:**
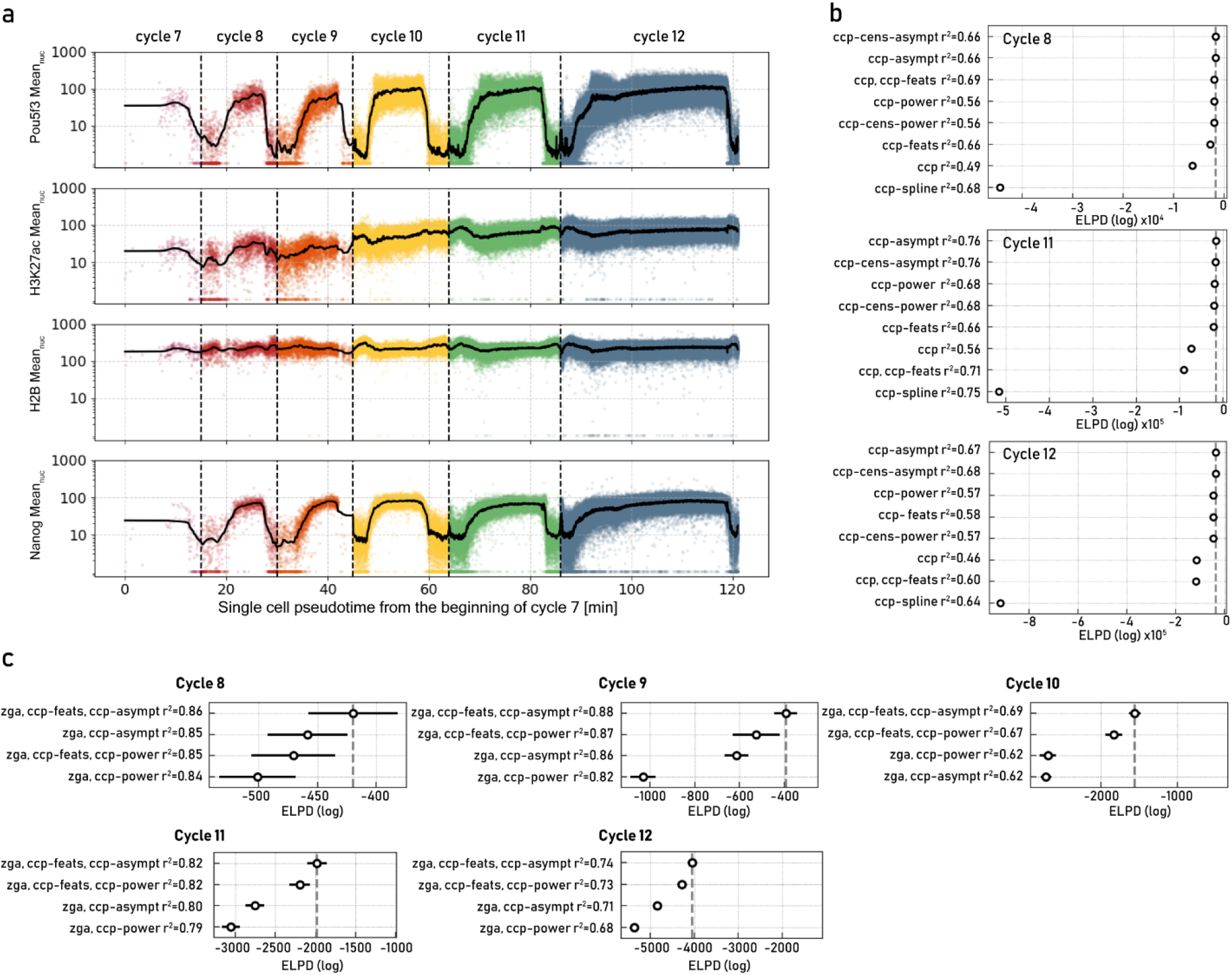
Modeling of transcriptional activity. a. Mean nuclear intensities of Pou5f3, H3K27ac, H2B and Nanog across division cycles 7 to 12. Sold lines demarcate Gaussian smoothing of points and dashed lines and color changes indicate cycle transitions. b. Model comparisons using CCP or features used for inferring CCP (CCP features, see Materials and Methods) as linear or non-linear predictors of transcription output measured by mean nuclear Pol-II-S2p (natural log-transformed) for cycles 8, 11 and 12. Models are ranked based on their performance on held-out data using expected log pointwise predictive density ELPD (improving from left to right). R^2^ demarcates explained variance for each model. Asymptotic (asympt), power and spline non-linear regression models of CCP were tested. Linear models include CCP or features used for CCP inference (ccp-feats). For the asymptotic and power regressions, additional models were tested where data was clipped at 100 gray values, censoring data points with low signal-to-noise ratios (ccp-cens-asympt, ccp-cens-power, Materials and Methods). c. Top 4 models in predicting mean nuclear Pol-II-S2p (natural log-transformed) using CCP as non-linear predictors and features used for inferring CCP (CCP features) and ZGA factors (in short zga including pou5f3, Nanog, H3K27ac and H2B) as linear regressor in cycles 8 to 12.

